# An ancestral informative marker panel design for individual ancestry estimation of Hispanic population using whole exome sequencing data

**DOI:** 10.1101/654939

**Authors:** Li-Ju Wang, Catherine W. Zhang, Sophia C. Su, Hung-I H. Chen, Yu-Chiao Chiu, Zhao Lai, Hakim Bouamar, Amelie G. Ramirez, Francisco G. Cigarroa, Lu-Zhe Sun, Yidong Chen

## Abstract

**Background:** Europeans and American Indians were major genetic ancestry of Hispanics in the U.S. In those ancestral groups, it has markedly different incidence rates and outcomes in many types of cancers. Therefore, the genetic admixture may cause biased genetic association study with cancer susceptibility variants specifically in Hispanics. The incidence rate and genetic mutational pattern of liver cancer have been shown substantial disparity between Hispanic, Asian and non-Hispanic white populations. Currently, ancestry informative marker (AIM) panels have been widely utilized with up to a few hundred ancestry-informative single nucleotide polymorphisms (SNPs) to infer ancestry admixture. Notably, current available AIMs are predominantly located in intron and intergenic regions, while the whole exome sequencing (WES) protocols commonly used in translational research and clinical practice do not contain these markers, thus, the challenge to accurately determine a patient’s admixture proportion without subject to additional DNA testing.

**Methods:** Here we designed a bioinformatics pipeline to obtain an AIM panel. The panel infers 3-way genetic admixture from three distinct continental populations (African (AFR), European (EUR), and East Asian (EAS)) constraint within evolutionary-conserved exome regions. Briefly, we extract ∼1 million exonic SNPs from all individuals of three populations in the 1000 Genomes Project. Then, the SNPs were trimmed by their linkage disequilibrium (LD), restricted to biallelic variants only, and assembled as an AIM panel with the top ancestral informativeness statistics based on the *I*_*n*_-statistic. The selected AIM panel was applied to training dataset and clinical dataset. Finally, The ancestral proportions of each individual was estimated by STRUCTURE.

**Results:** In this study, the optimally selected AIM panel with 250 markers, or the UT-AIM250 panel, was performed with better accuracy as one of the published AIM panels when we tested with 3 ancestral populations (Accuracy: 0.995 ± 0.012 for AFR, 0.997 ± 0.007 for EUR, and 0.994 ± 0.012 for EAS). We demonstrated the utility of UT-AIM250 panel on the admixed American (AMR) of the 1000 Genomes Project and obtained similar results (AFR: 0.085 ± 0.098; EUR: 0.665 ± 0.182; and EAS 0.250 ± 0.205) to previously published AIM panels (Phillips-AIM34: AFR: 0.096 ± 0.127, EUR: 0.575 ± 0.29; and EAS: 0.330 ± 0.315; Wei-AIM278: AFR: 0.070 ± 0.096, EUR: 0.537 ± 0.267, and EAS: 0.393 ± 0.300) with no significant difference (Pearson correlation, *P* < 10^-50^, *n* = 347 samples). Subsequently, we applied UT-AIM250 panel to clinical datasets of self-reported Hispanic patients in South Texas with hepatocellular carcinoma (26 patients). Our estimated admixture proportions from adjacent non-cancer liver tissue data of Hispanics in South Texas is (AFR: 0.065 ± 0.043; EUR: 0.594 ± 0.150; and EAS: 0.341 ± 0.160), with smaller variation due to its unique Texan/Mexican American population in South Texas. Similar admixture proportion from the corresponding tumor tissue we also obtained. In addition, we estimated admixture proportions of entire TCGA-LIHC samples (376 patients) using UT-AIM250 panel. We demonstrated that our AIM panel estimate consistent admixture proportions from DNAs derived from tumor and normal tissues, and 2 possible incorrect reported race/ethnicity, and/or provide race/ethnicity determination if necessary.

**Conclusions:** Taken together, we demonstrated the feasibility of using evolutionary-conserved exome regions to distinguish genetic ancestry descendants based on 3 continental-ancestry proportion, provided a robust and reliable control for sample collection or patient stratification for genetic analysis. R implementation of UT-AIM250 is available at https://github.com/chenlabgccri/UT-AIM250.

## Background

Over the past several hundred years, the America continent has been the hot spot to attracting peoples from difference continental populations that were originally separated by geography, such as African (mass immigrate by Atlantic slave trade), European (the age of exploration, and Spanish colonization of the Americas), and Asian (California gold rush) [1]. Due to meeting and mixing of previously isolated populations, through the years, the resulting *population admixture* carries novel genotypes with new genetic variations inherited DNA derived from a variety of ancestral populations [2], or in other words, admixed individuals have a genetic mosaic of ancestry that distinguishes them from their parental populations.

Hispanics in the U.S. have genetic ancestry from European, African and Native American. These admixture population present opportunity for the study of health disparity due to disease susceptibility [3, 4] or drug response [5–7]. In cancer study, it has been shown Hispanics are clearly different in many cancer incidence rates and outcomes [8]. However, the pattern of genetics and DNA variation of Hispanic individuals was affected by many historical events [9]. Therefore, genetic admixture may bias estimates of associations between candidate cancer susceptibility genes in Hispanics. The investigation of population structure and admixture proportion are more important in disease diagnosis. For example, the incidence rate and mutational frequency of liver cancer have been shown to be very different between Hispanic/Asian and non-Hispanic white populations [10], and especially Hispanic population in South Texas [11, 12]. To estimate the admixture proportion of individuals, most of published ancestry informative marker (AIM) panels were designed with up to a few hundred genome-wide ancestry-informative single nucleotide polymorphisms (SNPs) that exhibit large variation in minor allele frequency (MAF) among populations, and they are usually located in non-exonic regions [13–16]. The best AIM has the greatest difference in MAF between populations, they are also the best marker to determine admixture proportions. To estimate the admixture proportion, several model-based clustering approaches have been developed for the determination of the genetic ancestry of humans and other organisms. Pritchard et al. used a Bayesian algorithm STRUCTURE to first define the populations and then assign individuals to them [17]. A faster implementation algorithms ADMIXTURE was incorporated with a similar Bayes inference model, which allow the algorithm to be efficient for AIMs with more than thousands of markers [18]. More algorithms for estimating genetic ancestry can be found in literature [19].

Recently, whole exome sequencing (WES) has become the necessary strategy in translational research and clinical diagnostics to identify the underlying genetic cause of disease due to the fact that most of pathogenic variants are located in exonic regions, and also partially attributed to the drastically reduced cost of WES [20–22]. WES is a common protocol that provides detailed genetic variants including rare genetic events and unknown somatic mutations between different genetic conditions for large cohort of patients. Particularly in translational research, WES offers an unbiased view than conventional targeted molecular diagnostics approach, commonly available in many of large genomic studies such as genomic data at the Cancer Genome Atlas (TCGA) [23]. Previous studies showed that admixture proportions could be determined by using principal component analysis (PCA) with all variants [24], or using allele frequency for pooled DNA [25], and using off-target sequence reads [26]. However, using a panel of AIM within exome regions, if feasible, will allow rapid determination of a patient’s ancestry admixture from sequencing data and thus validate self-reported race/ethnicity categories.

In this study, we aimed to re-tune an AIM design pipeline to precisely characterize ancestry admixture of Hispanic populations using conventional WES data. Using 1000 Genomes Project data, we selected SNPs that have different MAF of African (AFR), European (EUR), and East Asian (EAS) populations by *I*_*n*_-statistics. We validated our optimal panel with 250 AIMs with the admixed American (AMR) of the 1000 Genomes Project, and compared our results to those from published AIM panels with SNPs optimally designed in intronic/intergenic regions. Finally, we demonstrated our AIM panel to TCGA-LIHC data and a Hepatocellular Carcinoma (HCC) study with self-reported Hispanic patients only enrolled in South Texas.

## Methods

### Population Samples

We use 1000 Genomes Phase III Whole Genome Sequencing data as the resource to calculate the ancestry informative markers (AIMs) [27]. In our study, the whole genome sequencing data was downloaded for each chromosome, excluding Mitochondrial, chrY, and chrX [ftp://ftp.1000genomes.ebi.ac.uk/vol1/ftp/]. 1000 Genome Phase III data is aligned with hg19 human reference genome. 1000 Genome SNPs were then extracted by ancestral populations (**Table 1**) using VCFtools [28] and BCFtools [29]. For the sake of simplicity, individuals from the Caribbean and African Americans were excluded from the ancestral population of Africa due to high levels of admixture observed. The Vietnamese population was also excluded from the East Asian ancestral population. Additionally, in order to eliminate Hispanics white interference, we pruned the Iberian population in Spain from the European population. For validation purpose, we utilized entire admixed American (AMR) collection, including Mexican Ancestry from LA, Puerto Ricans, Colombians and Peruvians (**Table 1**) to validate our panel.

**Table 1.** Populations of the 1000 Genome Project included in this study.

| Super Population | Subpopulation | # of samples |
| --- | --- | --- |
| **East Asian (EAS)** | Chinese Dai in Xishuangbanna (CDX),<br>Han Chinese (CHB),<br>Southern Han Chinese (CHS),<br>Japanese in Tokyo, Japan (JPT) | 405 |
| **African (AFR)** | Esan in Nigeria (ESN),<br>Gambian in Western Division, the Gambia (GWD),<br>Luhya in Webuye, Kenya (LWK),<br>Mende in Sierra Leone (MSL),<br>Yoruba in Ibadan, Nigeria (YRI) | 504 |
| **European (EUR)** | Utah residents (CEPH) with European Ancestry (CEU),<br>Finnish in Finland (FIN),<br>British in England and Scotland (GBR),<br>Toscani in Italia (TSI) | 396 |
| **Admixed American (AMR)** | Colombian in Medellin, Colombia (CLM),<br>Mexican Ancestry in Los Angeles, California (MXL),<br>Peruvian in Lima, Peru (PEL),<br>Puerto Rican in Puerto Rico (PUR) | 347 |

The populations were pulled from the 1000 genomes project database. We removed Vietnamese from EAS, African American from AFR, and Iberian of Spain from EUR (see Methods section).

### Data processing and Ancestry Informative Markers (AIMs) selection

The genome-wide data from the 1000 Genomes Project were first constrained to exonic region. Obtained SNPs were further subject to linkage disequilibrium filtering (*r*^2^ < 0.2, `plink option: --r2`), allele frequency (AF) calculation, and minor allele frequency (MAF < 0.01, `plink option: --maf 0.01`) elimination by PLINK (using vcftools to convert all three ancestral populations to .ped format with option `--plink`). The output files from PLINK were processed by the AIM generator (python script, AIMs_generator.py) [30]. This python script, provided by Daya *et. al*, performs LD pruning and select AIMs based on Rosenberg’s *I*_*n*_ Statistic [31] which defines the informativeness of SNPs,

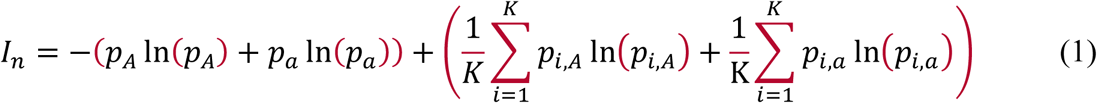

where *p*_*A*_ and *p*_*a*_ are the frequencies of 2 alleles, respectively, across all the individuals for a given marker, and *p*_*i,A*_, and *p*_*i,a*_ are the corresponding allele frequency in the *i*^th^ population. Notice that if a marker is unique in *i*^th^ population only, the second term in Eq. (1) will be 0, or *I*_*n*_ will be the largest, while *I*_*n*_ = 0 if the marker is equally distributed among all populations. To design our AIM panel, we first obtained nested subsets of AIMs up to 5000 SNP candidates (see **Additional Files 1: Table S1;** python code AIMs_generator.py, with ldfile/bim files from PLINK, ldthresh = 0.1, distances = 100000, strategy = *I*_*n*_), with min/max *I*_*n*_ for pair-wise ancestral populations as, AFR vs EUR: (0.000 to 0.614), AFR vs EAS: (0.000 to 0.623); and EAS vs EUR: (0.000 to 0.645), and the overall population (0.034 to 0.569). We expect 5000 SNP candidates will allow us to select robust AIM panel considering SNPs with balanced *I*_*n*_ from overall population, as well as least bias between pair-wise *I*_*n*_. The ancestry distribution of AIMs was provided in **Table 2**.

**Table 2.** Proportions of AIMs among 3 ancestral populations.

| # of AIMs | Populations |  |  |
| --- | --- | --- | --- |
|  | African | East Asian | European |
| 10 | 4 (40%) | 2 (20%) | 4 (40%) |
| 50 | 20 (40%) | 12 (24%) | 18 (36%) |
| 100 | 40 (40%) | 28 (28%) | 32 (32%) |
| 250 | 90 (36%) | 80 (32%) | 80 (32%) |
| 500 | 172 (34%) | 165 (33%) | 163 (33%) |
| 750 | 256 (34%) | 265 (35%) | 229 (31%) |
| 1000 | 329 (33%) | 355 (36%) | 316 (32%) |
| 2000 | 616 (31%) | 763 (38%) | 621 (31%) |
| 3000 | 920 (31%) | 1124 (38%) | 956 (32%) |
| 4000 | 1251 (31%) | 1488 (37%) | 1261 (32%) |
| 5000 | 1582 (32%) | 1810 (36%) | 1608 (32%) |

AIMs are determined by AIM\_generator.py script. We then examined AF of each population at each AIM location to assign the SNP to the dominate AF population (presented as the number of SNPs and percentage in each AIM panels). Note that AIM panels with more markers are not necessary contain less marker panels due to the requirement of balancing number of markers in 3 populations.

### Optimal AIM panel selection

The individual and population ancestral proportions were inferenced by STRUCTURE [17] and ADMIXTURE [18], as discussed in the Background section. The error of estimation of population proportions was determined by the result of STRUCTURE which were matched the race of corresponding individuals, or,

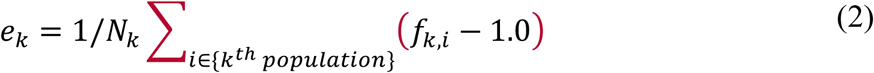

where we assume *f*_*k,i*_ is the admixture proportion of *i*^th^ person’s identified *k*^th^ population (ideally 100% in *k*^th^ population), and *k* = {EUR, ESA, and AFR}. A person will be classified into *k*^th^population if he/she has a maximum *k*^th^ population proportion estimated by STRUCTURE, thus we can estimate the error according to Eq. (2).

The optimal number of AIMs were determined when we observed accuracy, (1-*e*_*k*_), of classified known population does not improve by adding more SNP candidates from 5000 pools. We select AIMs with optimal balance in three populations (**Table 2**) with an equal number of SNPs from pair-wise *I*_*n*_ statistics. The final 250 AIMs (UT-AIM250) and its *I*_*n*_ Statistics were provided in **Additional Files 2: Table S2**.

### Whole Exome sequencing (WES)

The whole exome sequencing was performed with Illumina HiSeq 3000 system at GCCRI Genome Sequencing Facility, using Illumina’s TruSeq Rapid Exome Library Prep kit (Illumina, CA) which covers ∼45Mb with 99.45% of NCBI RefSeq regions. All exomeCapture sequencing was performed with 100bp paired-end (PE) module, and pooled the 6 samples per lane with targeted ∼100x fold coverage. Paired reads were aligned to human reference genome hg19 (exact the same of 1000 Genomes Project) with Burrows-Wheeler Aligner (BWA) [32]. Duplicated reads were removed by SAMtools [33] and Picard [http://broadinstitute.github.io/picard], and then realigned with GATK [34] considering dbSNPs information, and variants were identified by VarScan [35]. To report any variant statistics on locations specified by AIMs, we only require a minimum coverage threshold of 2, and no variant calling threshold.

### Principal Component Analysis (PCA)

PCA was performed on dataset of multi-locus genotypes to identify population distribution of each individual. The genotype matrix was obtained by applying the “read.vcfR” function of the R package [36]. Then, we converted the genotype to numeric numbers (0|0 = 0, 1|0 or 0|1 = 1, 1|1 = 2, and .|. = NA) by the Admixture_gt2PCAformat function (see github site). For PCA, we utilized dudi.pca (from “ade4” R package [37]). If there were missing value presence, we used estim_ncpPCA (form “missMDA” R package [38] to fill NA in genotype matrix) before performing PCA.

### AIM panel assessment

To assess the robustness of AIM panel that separates 3 continental populations, we first projected three populations into 3D space using PCA as described previously. We then assume each population possesses multi-variate Normal distribution,

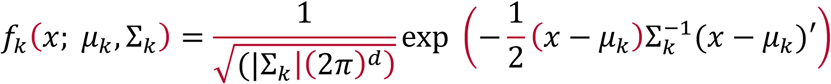

where *µ*_*k*_ is 1x*d* mean vector (here *d* = 3) of *k*^th^ population, and Σ_*k*_ is a *d*-by-*d* co-variance matrix. After estimation of the multivariate distributions of all 3 continental populations, we estimate the probability of mis-classify samples from one population to other two when the probability of a given sample with known population origin that has high probability to be assigned to the other two groups, or the misclassification probability of samples in *i*^th^ population into *j*^th^ population is We report the overall mis-classification probability, *P*_*m*_ (*i,j*) = ∫∫ _{*x* : *f i*(*x*) *< f j*(*x*)}_ *f* _*i*_ (*x* ; μ _*I*_, Σ _*i*_). *P* _*AIM*_ = Σ _all *I* ≠ *j*_ *P* _*m*_ (*i,j*) as a measure of the capacity separating populations using a specific AIM panel. The smaller *P*_AIM_ indicates less chance a sample to be misclassified using a given AIM panel, or in other words, farther separation between 3 populations.

### Hepatocellular Carcinoma (HCC) patients

We started by pruning in-house Whole Exome Sequencing data from 26 patients with Hepatocellular Carcinoma (HCC) with matched adjacent non-tumor and tumor. Initial pruning was performed by sequencing depth at each SNP, and further required biallelic SNPs (`vcftools options`: `--min-alleles 2 --max-alleles 2 --recode`). The SNPs were also eliminated if it had more than 10% missing genotype across all samples by VCFtools (`vcftools options: --max-missing 0.9 --recode`).

### The Cancer Genome Atlas (TCGA) – LIHC samples

We extracted specific SNP positions of UT-AIM250 from 788 TCGA-LIHC samples (376 patients) by using GDC BAM slicing tool (https://docs.gdc.cancer.gov/API/Users_Guide/BAM_Slicing/). The tool enables to download specific regions of BAM files instead of the whole BAM file for a given TCGA sample. These BAM slices were then processed with VarScan to determine variant fraction as described in previous sub-sections. The TCGA-LIHC whole exome data were derived from 4 sample types **(Figure 5A)**. According to race and ethnicity in clinical data of TCGA-LIHC, we re-classified 7 population groups (White, Asian, Black, Hispanic White, Reported as Hispanic, American Indian or Alaska Native, and Unknown) **(Figure 5A)**. The SNPs were conserved if it has more than 90% genotype throughout all sample by VCFtools, and further required biallelic SNPs.

## Results

### AIMs panel design and admixture estimation pipeline

We aim to design an AIM panel for estimating admixture proportion from the Hispanic population using conventional WES data. With this objective, we first focus our selection of continental population from the 1000 Genomes Project, removing all possible sources (removing African American from AFR collection and Iberian of Spain from EUR collection, and Vietnamese which are further down south of Asia) of error we can identify. We then constrain the ancestral markers in exome region. **Figure 1** outlined the flowchart of our AIM panel design pipeline (left panel). Here we assumed that our targeted population was comprised of three ancestry components: African (AFR), East Asian (EAS), and European (EUR). For this study, we focus only on SNPs (about 84.8 million variants total) that we extracted from three ancestry populations (*n* = 1,305) in 1000 Genomes (**Table 1**). These SNPs were then position filtered to ∼1 million exonic SNPs using VCFTools. To confirm these markers are good AIM candidate SNPs, all SNPs were pruned by those criteria: (1) linkage disequilibrium (LD) *r*^2^ < 0.2 within 100kb window to avoid the redundancy, (2) minor allele frequency (MAF) < 0.01 most likely are due to sequencing artifact, and (3) then the evaluation ancestral informativeness by using Eq. (1) *I*_*n*_-statistic value for all pair-wise comparison of 3 continental populations as described in the Methods Section. Total of 100,295 SNPs met the 1 and 2 criteria, and among them, we generated AIMs panels with 10, 50, 100, 250, 500, up to 5000 AIMs (see **Table 2**, and **Additional Files 1: Table S1**). For AIM 250, we have min/max *I*_*n*_ for pair-wise ancestral populations from 250 markers: AFR vs EUR: (0.000 to 0.614), AFR vs EAS: (1.185×10^-5^ to 0.623); and EAS vs EUR: (0.000 to 0.645), and overall population (0.134 to 0.569) (**Additional Files 2: Table S2).**

**Figure 1.**
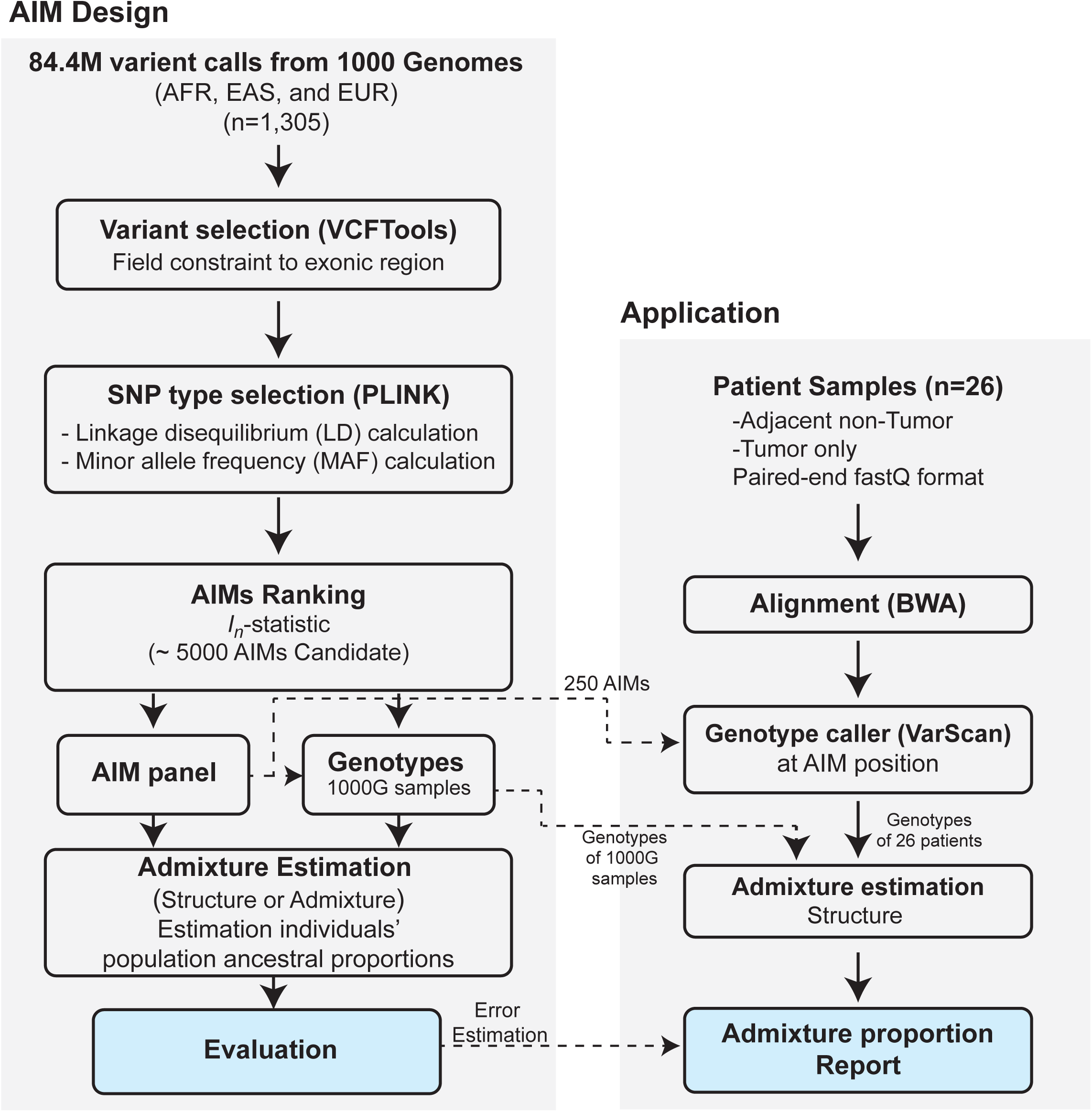
Flowchart of our AIM panel design and analysis pipelines. The pipeline is separated into two parts, AIM panel design (AIM Design) and Ancestral proportion estimation application (Application). For AIM Design pipeline (left panel), Variant files form the 1000 Genome Project (*n* = 1,305) were position filtered to exonic region by VCFTools. The variant files were calculated linkage disequilibrium (LD) and minor allele frequency (MAF) by PLINK. SNPs were selected as ancestry informativeness markers base on *I*_*n*_-statistic for overall population or each continental population. Finally, population ancestral proportions were estimated by STRUCTURE. For the Application (right panel), the 26 HCC tumors with matched adjacent non-tumor data were processed by standard whole exome sequencing analysis pipeline using BWA, GATK and then genotype caller VarScan at AIM positions. The last step in this panel was admixture estimation and reported the ancestral proportions of individual.

### Comparison of population structure tools

Here we compared the two popular admixture tools, STRUCTURE and ADMIXTURE. These two tools utilized different algorithms (Bayesian statistics vs maximum likelihood estimation) to estimate population structure. In addition, the execution time of ADMIXTURE was much more efficient than STRUCTURE without much sacrifice in accuracy, including multiple threads setting in ADMIXTURE. As expected, the accuracy of population estimation between both tools showed that STRUCTURE was better than ADMIXTURE (both set at *K* = 3) (**Figure 2A, B**). For each population and its corresponding ancestral proportion estimation, the mean and standard deviation (SD) of ancestry estimation accuracy between STRUCTURE and ADMIXTURE was AFR: 0.991 ± 0.016 vs 0.977 ± 0.027 (*P* = 7.20×10^-23^, *t*-test, one-tails), EUR: 0.988 ± 0.021 vs 0.969 ± 0.034 (*P* = 1.70×10^-20^), and EAS: 0.996 ± 0.009 vs 0.989 ± 0.017, (*P* = 2.92×10^-13^). At 250 AIMs, we observed the best grouping accuracy and lowest SD in three ancestral populations with STRUCTURE algorithm (AFR: 0.995 ± 0.012, EUR: 0.994 ± 0.012, and EAS: 0.997 ± 0.007), while ADMIXTURE need more than 250 AIMs to gain better accuracy (**Figure 2A, B**). Examining individual’s estimation carefully from both algorithms further confirmed that ADMIXTURE is less robust on estimation (**Figure 2C, D**, and much longer green tail in **Figure 2D**, inset for the AFR population). For these reasons, subsequent analysis was mostly focused on the AIM 250 panel (termed UT-AIM250 thereafter) and the STRUCTURE algorithm for admixture proportion estimation. Within UT-AIM250 panel, we identified 90 African AIMs (36%), 80 European AIMs (32%), and 80 East Asian AIMs (32%) (see **Table 2** and **Additional Files 2: Table S2**). We utilized genotypes from three ancestry populations (*n* = 1,305) in 1000 Genomes on UT-AIM250 panel and confirmed that the UT-AIM250 has sufficient discriminating capacity to separate three ancestral populations, as shown in principal component plot (**Figure 2E**, with 95% and 99% confidence circles in solid and dash lines, respectively).

**Figure 2.**
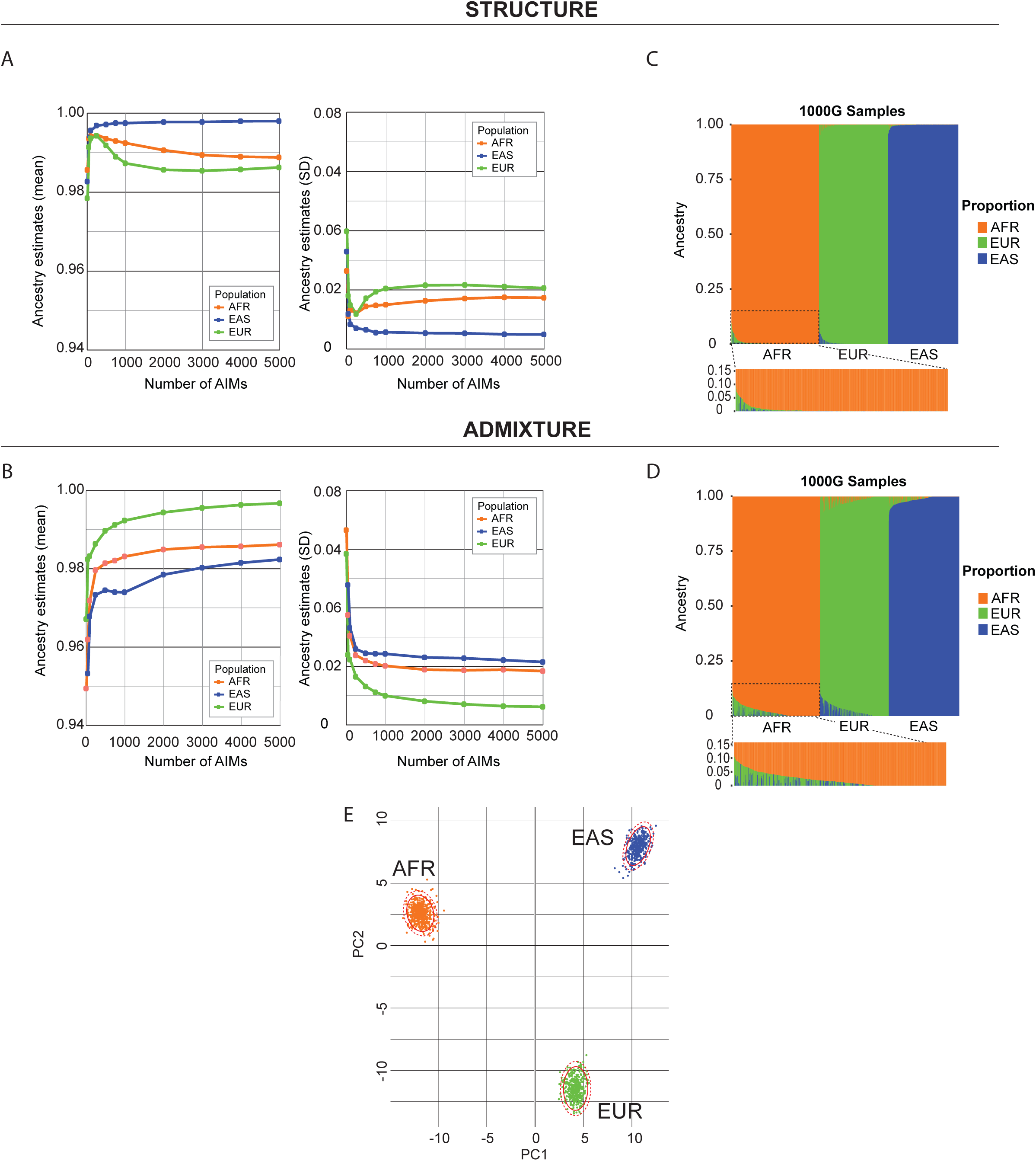
Selection of a tool for ancestral population proportion estimation. The results were presented as those from STRUCTURE (A, C) and from ADMIXTURE (B, D). **(A, B)** Performance of AIM panels with different number of markers. Mean and Standard deviation were plotted for each population. At 250 markers, the accuracy plateaus when STRUCTURE algorithm is used. **(C, D)** Proportion plot for ancestral populations on 250AIMs between STRUCTURE and ADMIXTURE. The populations were ordered by groups: AFR: African, EUR: European, and EAS: East Asian. Individuals in (D) were ordered according to (C). We also zoomed the part of AFR out to further demonstrate the population distribution using two different algorithms. **(E)** Principal component analysis (PCA) plot for three ancestral population distribution on 250 AIMs.

### Comparison between our UT-AIM250 panel and published 34-AIM and 278-AIM panels

To further examine the accuracy, we compared our UT-AIM250 panel and two published panels, 34 AIM-panel [14] (termed Phillips-AIM34) and 278 AIM-panel [39] (termed Wei-AIM278), on the Admixed American (AMR) population also from the 1000 Genomes Project. Those panels were generated from three continental populations (with slightly inclusion criterion and samples available at the time): African (AFR), European (EUR), and East Asian (EAS). The differences between panels are the position of AIMs and AIM selection approach: the Phillips-AIM34 panel is composed of SNPs in exonic regions (2 SNPs) and non-exonic regions (32 SNPs); the Wei-AIM278 panel is composed of SNPs in exonic (3 SNPs) and non-exonic regions (275 SNPs). **Figure 3** depicts the results from UT-AIM250 (**Figure 3A, B**), Phillips-AIM34 (**Figure 3C, D**) and Wei-AIM278 panels (**Figure 3E, F**) of 3 continental ancestral populations plus Admixed American (AMR). The AMR was composed of four subpopulations, Colombian (CLM), Mexican in LA (MXL), Peruvian (PEL), and Puerto Rican (PUR). Following the analysis pipeline (**Figure 1**, right panel), the genotypes of AMR (*n* = 347) was extracted by according to UT-AIM250, Phillips-AIM34, and Wei-AIM278, and then merged with genotypes from 3 continental populations (*n* = 1,305, for a total of 1,652 individuals). The structure of populations was estimated by STRUCTURE and plot by both bar-chart and principal components (**Figure 3**). As shown in the figure, three panels can separate continental populations, and UT-AIM250 provided a much superior population separation (**Figure 3A, C, E**), with misclassification probability *P*_UT-AIM250_, *P*_Phillips-AIM34_, and *P*_Wei-AIM278_ to be 4.563×10^-37^, 2.059 ×10^-5^, and 3.221 ×10^-26^, respectively (see Methods section). The resulted population structure showed a similar trend of three panels (**Figure 3B, D, F**): within AMR sub-populations, Puerto Rican had much strong European ancestral proportion (AFR : 0.149 ± 0.109, EUR : 0.789 ± 0.111, and EAS : 0.062 ± 0.051), while Peruvian had the strongest influence from Asian (AFR : 0.032 ± 0.066, EUR : 0.449 ± 0.111 and EAS : 0.519 ± 0.124), in line with previous published studies [13, 40, 41]. In terms of MXL, the proportions for 3 ancestral populations were AFR = 0.046 ± 0.046, EUR = 0.634 ± 0.142, and EAS = 0.320 ± 0.149. Although UT-AIM250, Phillips-AIM34, and Wei-AIM278 showed different distribution of ∼1,600 individuals, Pearson correlation between three panels confirmed their high similarity (**Table 3**). We also examined their proportion obtained from three panels to perform AFR-AFR correlation over all AMR individuals. The correlation ρ were 0.70, 0.83 and 0.85 (Phillips-AIM34); 0.89, 0.93 and 0.96 (Wei-AIM278) for AFR, EUR and EAS ancestral proportion, respectively. Similar correlation coefficients for each sub-population can be found in **Table 3**.

**Table 3.** Pearson correlation coefficients between UT-AIM250 and published panels of the AMR population.

| Panel | Population | Pearson Correlation (p-value) |  |  |  |  |
| --- | --- | --- | --- | --- | --- | --- |
|  |  | PUR<br>(n = 104) | CLM<br>(n = 94) | PEL<br>(n = 85) | MXL<br>(n = 64) | All<br>(n = 347) |
| Phillips-AIM34 | AFR-AFR $\rho$ | 0.67<br>($P = 5.97 \times 10^{-15}$) | 0.57<br>($P = 2.11 \times 10^{-9}$) | 0.83<br>($P = 1.46 \times 10^{-22}$) | 0.22<br>($P = 7.56 \times 10^{-2}$) | 0.70<br>($P = 5.83 \times 10^{-52}$) |
| | EUR-EUR $\rho$ | 0.69<br>($P = 9.72 \times 10^{-16}$) | 0.67<br>($P = 9.39 \times 10^{-14}$) | 0.57<br>($P = 1.60 \times 10^{-8}$) | 0.81<br>($P = 4.09 \times 10^{-16}$) | 0.83<br>($P = 7.30 \times 10^{-91}$) |
| | EAS-EAS $\rho$ | 0.26<br>($P = 6.87 \times 10^{-3}$) | 0.42<br>($P = 2.00 \times 10^{-5}$) | 0.64<br>($P = 4.18 \times 10^{-11}$) | 0.77<br>($P = 1.49 \times 10^{-13}$) | 0.85<br>($P = 4.75 \times 10^{-96}$) |
| Wei-AIM278 | AFR-AFR $\rho$ | 0.89<br>($P = 1.96 \times 10^{-37}$) | 0.86<br>($P = 1.75 \times 10^{-28}$) | 0.89<br>($P = 2.29 \times 10^{-30}$) | 0.40<br>($P = 9.75 \times 10^{-4}$) | 0.89<br>($P = 1.20 \times 10^{-122}$) |
| | EUR-EUR $\rho$ | 0.80<br>($P = 1.74 \times 10^{-24}$) | 0.84<br>($P = 1.55 \times 10^{-26}$) | 0.83<br>($P = 6.24 \times 10^{-23}$) | 0.92<br>($P = 1.24 \times 10^{-26}$) | 0.93<br>($P = 1.60 \times 10^{-152}$) |
| | EAS-EAS $\rho$ | 0.47<br>($P = 3.89 \times 10^{-7}$) | 0.73<br>($P = 7.51 \times 10^{-17}$) | 0.89<br>($P = 1.01 \times 10^{-29}$) | 0.93<br>($P = 8.98 \times 10^{-29}$) | 0.96<br>($P = 6.04 \times 10^{-193}$) |

Lower correlation coefficient is expected if proportion of certain ancestral population is smaller than its estimation variation.

**Figure 3.**
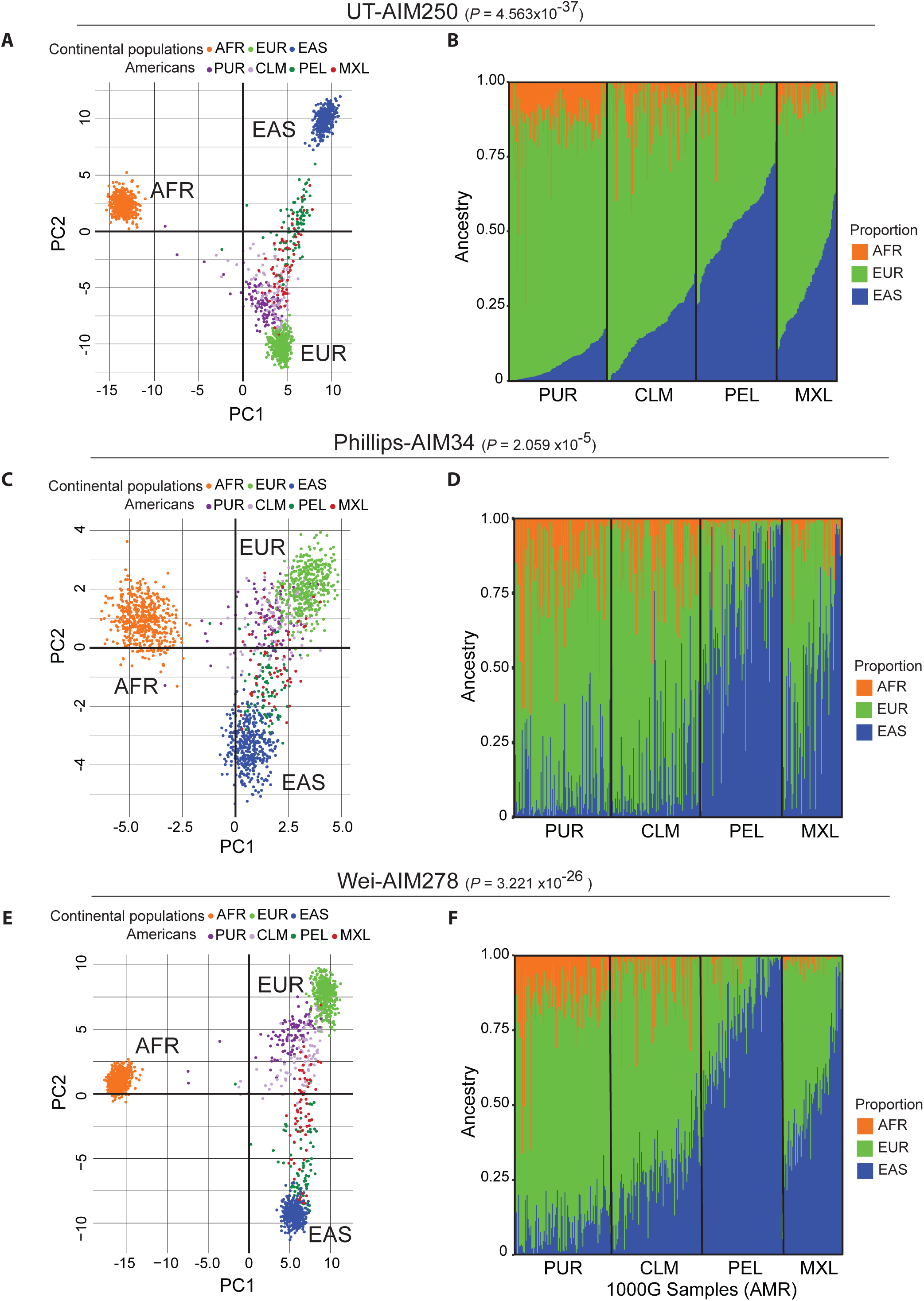
Comparison between the proposed UT-AIM250 panel and two published AIM panels. (A, C, E) Principal component analysis (PCA) plot for AMR population distribution on UT-AIM250, Phillips-AIM34, and Wei-AIM278 panels. **(B, D, F)** Proportion plot for admixed Americans (AMR). The population were arranged by individuals within groups: PUR: Puerto Rican; CLM: Colombian; PEL: Peruvian and MXL: Mexican in LA.

### Ancestry estimation for HCC patients

The key to design UT-AIM250 is to validate self-reported Hispanic patients for translational study without adding specific ancestral markers to the standard Exome Capture kit for sequencing library preparation. We applied UT-AIM250 panel to estimate the ancestral proportion of a collection of 26 HCC patients with matched tumor tissues and adjacent non-tumor (Adj. NT) tissues. We extracted genotypes at 250 SNPs locations from Adj. NT and tumors using VarScan (see Methods), merged with 1,305 continental populations from 1000 Genomes Project, and visualized using first 2 principal components (**Figure 4A** for Adj. NT and **4B** for tumor only). No obvious differences were observed between Adj. NT and tumor samples. They were located between the EAS and EUR clusters, indicating the possibility of using tumor data alone to assess the patient ancestral proportion. In addition, we calculated ancestral component by STRUCTURE (*K* = 3). The ancestral proportions of our HCC patients (all self-reported as Hispanic from San Antonio or South Texas regions) are AFR = 0.065 ± 0.044, EUR = 0.595 ± 0.151, and EAS = 0.340 ± 0.163, similar to those of MXL. In the triangle plots (**Figure 4C, E)**, HCC patients mostly aligned along the axis of EAS and EUR, similar to the PCA plot. One patient (HCC-3) was predicted as Asian (in the Asian population in PCA plot, and Asian proportion = 0.916; **Figure 4D, F**), so we excluded this patient from subsequent genetic analysis. Similar to the comparison between STRUCTURE and ADMIXTURE algorithm, we examined the correlation coefficient ρ between tumor tissues and Adj. NT tissues. The results were 0.96, 0.99 and 0.99 for AFR-AFR, EUR-EUR, and EAS-EAS, respectively (all *P* < 10^-14^). Taken together, our UT-AIM250 panel is accurate and robust to determine the ancestral proportion from normal or even tumor samples.

**Figure 4.**
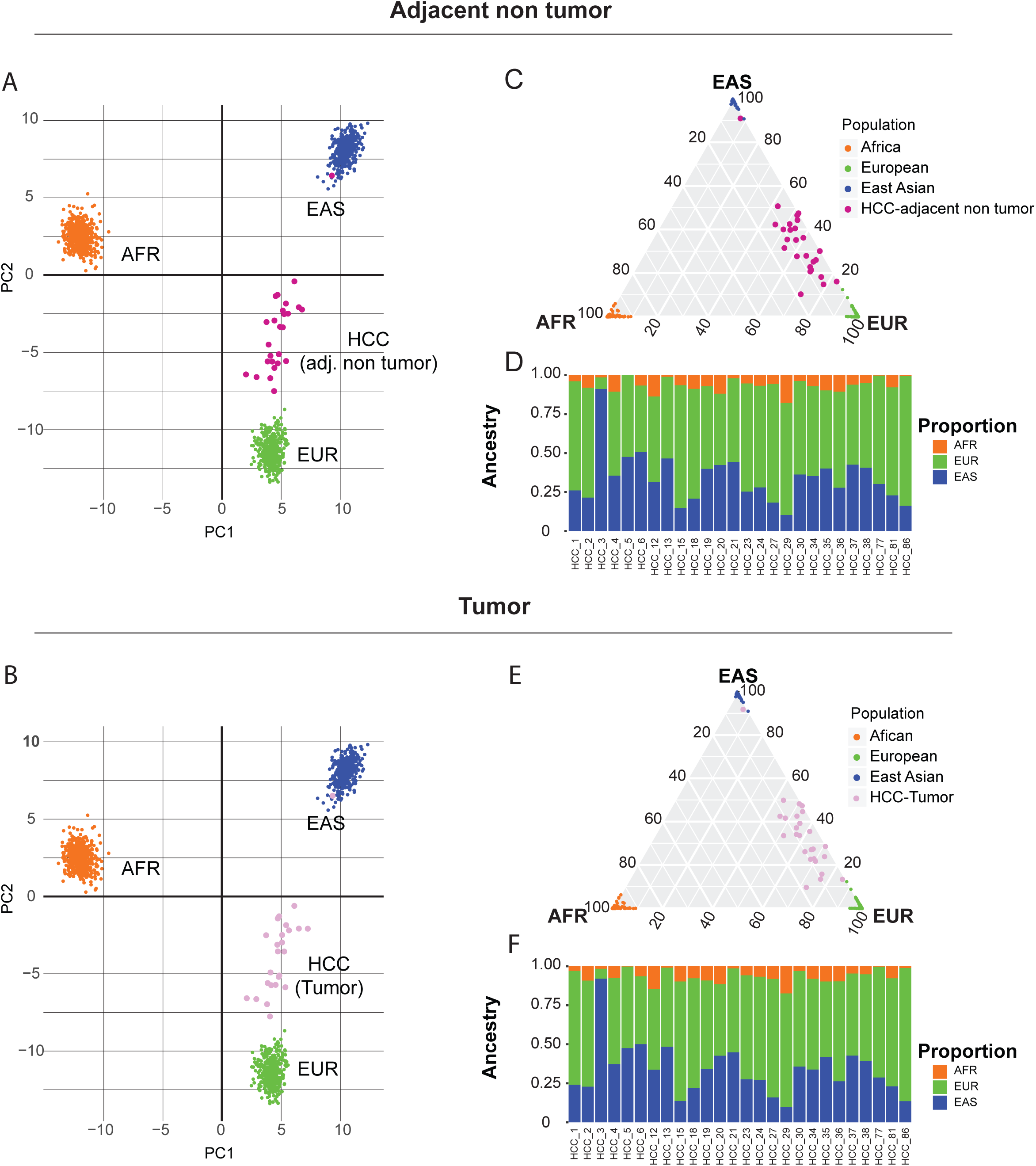
Application to 26 HCC tumors with matched adjacent non-tumor using exome-seq data. (A, B) PCA plots from HCC adjacent non-tumor samples and HCC tumor samples. **(C, E)** Triangle plot for ancestral division probability of HCCs from African (AFR), East Asian (EAS), and European (EUR). **(D, F)** The proportion plot for HCCs, ordered by the number of individuals.

**Figure 5.**
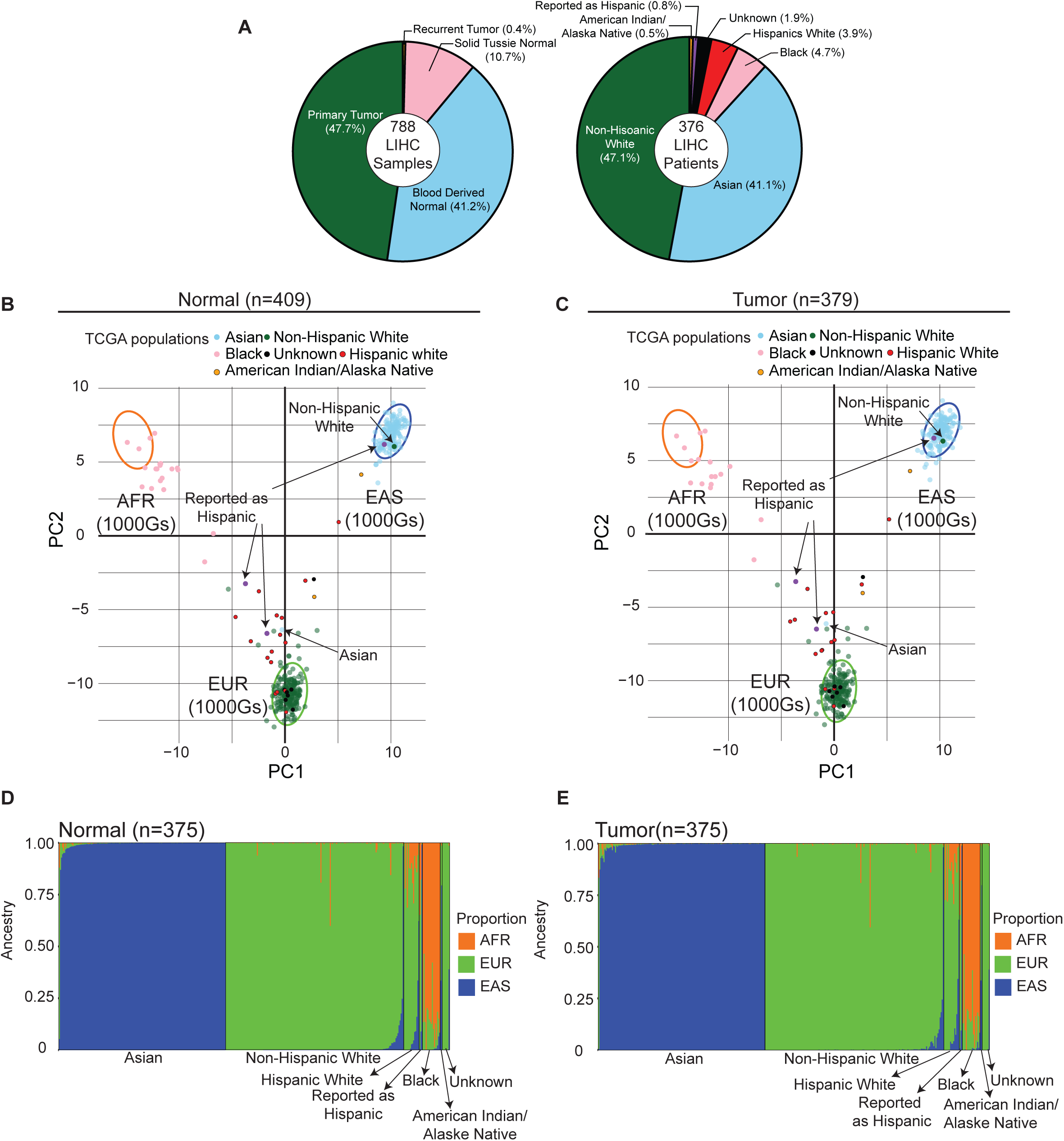
Application to 788 TCGA-LIHC samples (376 patients). **(A)** Summary of TCGA-LIHC samples and patients. Left pie-chart is the break-down of sample types, and right pie-chart is the break-down of self-reported race and ethnicity of all LIHC patients. **(B, C)** PCA plots from 788 TCGA-LIHC samples (Normal: *n* = 409; Tumor: *n* = 379). Normal group includes DNAs derived from blood and/or solid tissue normal, and tumor group includes primary tumor and recurrent tumor. Purple points were these from patients “Reported as Hispanic”. The confidence interval depicted by three ellipses (determined from 3 continental population EAS, EUR and AFR of 1000 Genome Project) is 0.99. **(D, E)** The proportion plot for 375 TCGA-LIHC patients who matched normal (Blood derived normal: *n* = 325; Solid tissue normal: *n* = 50) and primary tumor samples, ordered by the number of individuals.

### Ancestry estimation for TCGA-LIHC samples

In order to verify the accuracy of UT-AIM250 on different sample types, we evaluated all TCGA-LIHC 376 patients and compared to their self-reported race/ethnicity. TCGA-LIHC has total of 788 whole exome sequencing (WES) data, derived from 4 sample types (41.2% Blood Derived Normal, 10.7% Solid Tissue Normal, 47.7% Primary Tumor, and 0.4% Recurrent Tumor, **Figure 5A** left panel, and **Table S3)**. Based on race and ethnicity of each patient reported, we divided all 376 patients to 7 populations (47.1% White, 41.1% Asian, 4.7% Black, 3.9% Hispanic white, 0.8% Reported as Hispanic, 0.5% American Indian or Alaska Native, and 1.9% Unknown, **Figure 5A** right panel, and **Table S3**). We applied UT-AIM250 to all 788 samples (normal *n* = 409, and tumor *n* = 379). The PCA plots showed similar patterns in both normal and tumor (**Figure 5B** for normal and **5C** for tumor only), indicating our UT-AIM250 panel is robust even if normal DNA is not available. In **Figure 5B-E**, we selected 375 TCGA-LIHC patients with matched primary tumor samples and normal samples (325 Blood Derived normal and 50 Solid tissue normal), excluding TCGA-BC-4072 which has primary tumor sample only. We utilized STRUCTURE (K = 3) to calculate ancestral components (**Table S3**). The ancestral proportions of 375 TCGA-LIHC patients were plotted with bar-chart (**Figure 5D** for normal, **5E** for primary tumor), ordered by the number of individual patients. Among 375 TCGA-LIHC patients, there are two patients, TCGA-DD-AACA and TCGA-ZS-A9CF, have three sample types, Blood Derived Normal, Primary Tumor, and Recurrent Tumor. We compared the ancestral proportions of three sample types on each person, and the results were consistent (TCGA-DD-AAC: EAS = 0.999, EUR = 0.001, and AFR = 0; TCGA-ZS-A9CF: EAS = 0.001, EUR = 0.999, and AFR = 0.001). Our analysis also concluded that there were three patients (TCGA-G3-A5SI, TCGA-G3-AAUZ, and TCGA-FV-A4ZQ) who do not match the race and ethnicity from their self-reported race/ethnicity status. TCGA-G3-A5SI (self-reported as Asian) was predicted as white (EUR proportions = 0.826; **Figure 5B-C**). We also predict both patients TCGA-G3-AAUZ (self-reported as Hispanics) and TCGA-FV-A4ZQ (self-reported as White) to be Asian (EAS proportion = 0.992, and 0.984, respectively). In addition, 7 patients with unknown race/ethnicity status were assigned to their corresponding genetic groups. Therefore, the SNP positions of our UT-AIM25 is unaffected by possible tumor mutations and UT-AIM250 is a robust panel of ancestral markers within exome regions.

## Discussion

In this study, we developed, validated and tested the pipeline for designing AIM panels within the evolutionary-conserved exome regions to distinguish genetic ancestry descendants base on three continental populations (African, European, and East Asian). Although WES could be applied to analyze population structure using all variants [24], it will be problematic since variants will be influenced by the number of somatic mutations in tumor samples, which typically are significantly different on germline and tumor [42–44]. By using UT-AIM250 panel and we acquired the satisfactory result as we expected (by removing low-frequency MAFs and constraint with only biallelic SNPs) even with the tumor samples. To further reduced the impact of somatic mutations to our AIM panel design, one may choose to filter SNPs from COSMIC database [45] or other relevant tumor variant collections, such as the International Cancer Genome Consortium (ICGC) [46]. While the number of our HCC patients is small, we believe it is sufficient to demonstrate the utility of WES data to identify ancestry proportion of individuals. In some clinical applications in which only tumor samples are available, our UT-AIM250 is proved to be a cost-effective tool to confirm the race and ethnicity of patients when WES data are available.

The AIMs were selected from three continental populations (African, European, and East Asian). Those populations were the major groups which contributed to the ancestral genetic variety of people in the US through various migration routes, and some of them are extremely complex [47]. There are variable phenotypes of Hispanics in the US [48], and it is recognized that health disparity does exist in different populations in the US, but also within Hispanic populations [49] due to their diverse genetic background such as populations shown in **Figure 3A** (AMR subpopulations). We have carefully selected subpopulations specifically for our targeted population, such as removing of Iberian (IBS), to further constrain EUR to be considered as Non-Hispanic White (NHW).

Another important factor is to use the Native American as another continental population. However, as shown in **Figure 3A**, choose which subpopulation in AMR collection of the 1000 Genomes Project is a challenging task. We will continue evaluating the genomic resources, preferred whole genome sequencing (WGS) data of the Native American that are commonly accepted as an ancestral population. We chose Asian not only due to its stable genome variation, but also because of the convincing evidence that one of the origin ancestries of Native American could be Asian who came from northeast Asia by passing Beringia strait [50, 51]. We believe our AIM panel shall be sufficient for us to identify distinct genetic groups for downstream data analysis, such as risk factor assessment.

Along with the development of precision medicine, the population determination plays an important role [52]. Whether our HCC patients or TCGA-LIHC patients, they met the same problem about accuracy of patients self-reported race/ethnicity status in this study. After ancestral estimation, the results of some patients do not match reported. Due to several potentially factors, such as native language, environment, immigration, etc., patients usually caused misjudgment to their real population, especially on immigrant society. Base on this reason, UT-AIM250 could correct this mistake and provide confidential population report for medical treatment.

There are many different types of variants other than SNP, such as insertions, deletions, and haplotypes. In this study, we focused on biallelic SNPs. Extending to insertion and deletion will complicate the analysis due to the precise definition of the indel in each person. Furthermore, in the population genetic field, it was considered and analyzed other potentially factors on the distribution of population proportions [53]. Clearly, more works remain to be done in the future.

## Conclusions

We have constructed a unique AIM panel, UT-AIM250 which was designed within the evolutionary-conserved exome regions, to determine the admixture proportions of three continental populations (AFR, EUR, and EAS) for Hispanic in South Texas. To evaluate our panel, we demonstrated the accuracy using AMR subpopulations from the 1000 Genomes Project and compared the results obtained from the Phillips-AIM34 and Wei-AIM278 panels. We further applied our panel on 26 Hispanic HCC patients and 375 TCGA-LIHC patients with matched tumor and adjacent non-tumor tissues. The estimated ancestral proportions shown no significant difference between those obtained from non-tumor tissues and tumor tissues, enable us to evaluate patient specimen with tumor DNA only to verify self-reported Hispanic patients and/or their specific genetic analysis groups. Since WES is one of the dominant genome-wide variant analysis platforms, we UT-AIM250 panel, offers a cost-effective yet accurate method for the determination of patients’ ancestral composition. R implementation of UT-AIM250 is available at https://github.com/chenlabgccri/UT-AIM250.

## Abbreviations

Adj. NT: adjacent non-tumor
AFR: African
AIM: Ancestry Informative Marker
AMR: Admixed American
BWA: Burrows-Wheeler Aligner
CLM: Colombian
EAS: East Asian
EUR: European
HCC: Hepatocellular Carcinoma
LD: Linkage Disequilibrium
MAF: Minor Allele Frequency
MXL: Mexican in LA
PCA: Principal Component Analysis
PE: Paired-end
PEL: Peruvian
PUR: Puerto Rican
SD: Standard Deviation
SNPs: Single Nucleotide Polymorphisms
TCGA-LIHC: TCGA Liver Hepatocellular Carcinoma
TCGA: The Cancer Genome Atlas
WES: Whole Exome Sequencing
WGS: Whole Genome Sequencing

## Declarations

### Acknowledgments

The authors greatly appreciate all members of the research teams for HCC study at the UT Health Science Center at San Antonio for their inputs into the biology and statistical methods of this work. We also thank the patients who donated their tissues for the study. The TCGA-LIHC datasets used for the analyses described in this manuscript were obtained from dbGap at https://www.ncbi.nlm.nih.gov/gap/ through dbGap accession phs000178.v10.p8 under the South Texas Liver Cancer Study (project ID : 13485). The results published here are in whole or part based upon data generated by The Cancer Genome Atlas managed by the NCI and NHGRI. Information about TCGA can be found at https://cancergenome.nih.gov/.

### Availability of data and material

R implementation and example of UT-AIM250 is available at https://github.com/chenlabgccri/UT-AIM250.

### Authors’ contributions

All of the authors conceived the study. LJW, CZ and YC designed the pipeline. LJW, CZ, and SS performed data analysis on 1000 Genomes samples. HB, ZL, LZS and FGC provided HCC samples and data. YCC and HHC performed data preprocessing. LJW, YCC, LZS, AGR, FGC and YC interpreted the data. LJW and YC wrote the manuscript. All of the authors have read and approved the final manuscript.

### Ethics approval and consent to participate

Not applicable.

### Consent for publication

Not applicable.

## Additional files

**Additional files 1: Table S1.** Informativeness of AIMs across different panels.

**Additional files 2: Table S2.** Informativeness of AIMs of the UT-AIM250 panel.

**Additional files 3: Table S3A and B.** The ancestral proportions and clinical data of TCGA-LIHC samples (S3A tumor DNAs, S3B normal DNAs).

